# Successful mating and hybridisation in two closely related flatworm species despite significant differences in reproductive morphology and behaviour

**DOI:** 10.1101/851972

**Authors:** Pragya Singh, Daniel Ballmer, Max Laubscher, Lukas Schärer

## Abstract

Speciation is usually a gradual process, in which reproductive barriers between two species accumulate over time. Reproductive traits, like genital morphology and mating behaviour, are some of the fastest diverging characters and can serve as reproductive barriers. The free-living flatworm *Macrostomum lignano*, an established model for studying sex in hermaphrodites, and its congener *M. janickei* are closely related, but differ substantially in their male intromittent organ (stylet) morphology. Here, we examine whether these morphological differences are accompanied by differences in behavioural traits, and whether these could represent barriers to successful mating and hybridization between the two species. Our data shows that the two species differ in many aspects of their mating behaviour, with *M. janickei* having a five-fold longer copulation duration, copulating less frequently, and having a longer and more delayed suck behaviour (a postcopulatory behaviour likely involved in sexual conflict). Interestingly, and despite these significant morphological and behavioural differences, the two species mate readily with each other in heterospecific pairings, often showing behaviours of intermediate duration. Although both species have similar fecundity in conspecific pairings, the heterospecific pairings revealed clear postmating barriers, as only few heterospecific pairings produced F1 hybrids. These hybrids had a stylet morphology that was intermediate between that of the parental species, and they could successfully backcross to both parental species. Finally, in a mate choice experiment we tested if the worms preferentially mated with conspecifics over heterospecifics, since such a preference could represent a premating barrier. Interestingly, the experiment showed that the nearly two-fold higher mating rate of *M. lignano* caused it to mate more with conspecifics, leading to assortative mating, while *M. janickei* ended up mating more with heterospecifics. Thus, while the two species can hybridize, the mating rate differences could possibly lead to higher fitness costs for *M. janickei* compared to *M. lignano*.

## Introduction

The biological species concept defines species as groups of individuals that interbreed in nature to produce viable and fertile offspring (Mayr 1942; Coyne and Orr 2004). They are usually isolated from interbreeding with other species by reproductive barriers, though in some cases they remain capable of producing hybrid offspring with closely related species. Accordingly, an important step for the origin and maintenance of species is the evolution of reproductive barriers, which are usually split into prezygotic and postzygotic barriers (Butlin et al. 2012; Ostevik et al. 2016; Lackey and Boughman 2017; Sato et al. 2018). While prezygotic barriers involve the prevention of zygote formation, postzygotic barriers lead to zygote mortality, or inviable or sterile hybrid offspring that are unable to pass on their genes. Moreover, prezygotic barriers can be ecological, temporal, behavioural, mechanical or gametic, and can be further subdivided into premating barriers and postmating-prezygotic barriers. Premating barriers act to prevent the occurrence of heterospecific matings. For example, if a species has a mating preference for conspecific partners over heterospecifics, this mating preference can lead to assortative mating between conspecifics and thereby function as a premating barrier (Williams and Mendelson 2010; Ciccotto et al. 2013; Zhou et al. 2015). Postmating-prezygotic barriers often involve conspecific sperm precedence due to postcopulatory processes, such as sperm competition and cryptic female choice, or they can result from an incompatibility of female reproductive organs with heterospecific male ejaculate (Manier et al. 2013; Soudi et al. 2016; Firman et al. 2017; Devigili et al. 2018; Garlovsky and Snook 2018; Turissini et al. 2018). However, in studies of internally fertilizing species it can often be difficult to distinguish whether the barrier is prezygotic (e.g. if despite mating, heterospecific sperm is not transferred or lost in the female reproductive trait) or postzygotic (e.g. if any resulting zygotes do not develop properly).

Species in the early stages of divergence will often not have complete reproductive barriers between them, but as they diverge in their traits, more reproductive barriers usually accumulate over time, since these divergent traits can function as barriers. Reproductive traits may diverge particularly quickly, since they are the primary targets of sexual selection, often leading to rapid accumulation of phenotypic differences (Eberhard 1985; Arnqvist 1997; Swanson and Vacquier 2002; Gröning and Hochkirch 2008). Therefore, sexual selection can play an important role in evolutionary diversification, reproductive isolation and speciation (Kraaijeveld et al. 2011; Janicke et al. 2018 but see Morrow et al. 2003). This is supported by the fact that reproductive traits, such as mating behaviour and genital morphology, have been shown to diversify faster than other traits (Arnqvist 1998; Gleason and Ritchie 1998; Puniamoorthy et al. 2009, 2010; Puniamoorthy 2014) and can differ markedly even between recently diverged species (Schärer et al. n.d.; Anthes and Michiels 2007; Puniamoorthy et al. 2009, 2010; Kelly and Moore 2016), and sometimes even between populations of the same species (Herring and Verrell 1996; Klappert et al. 2007; Puniamoorthy 2014). Moreover, some studies have shown that mating behaviour might evolve even more quickly than genital morphology (Puniamoorthy 2014). Thus, a rapidly evolving reproductive trait like reproductive behaviour can represent a premating barrier by being involved in mate recognition and assortative mating (Herring and Verrell 1996; Ritchie et al. 1999), while a difference in genital morphology can prevent successful mating and thus represent a mechanical barrier (Masly 2012; Barnard et al. 2017).

In recently diverged species that occur in sympatry, selection may occur to reduce the likelihood of heterospecific reproductive interactions, whenever such interactions lower individual fitness (either directly or via low fitness hybrids). This selection can cause greater divergence in reproductive traits, leading to reproductive character displacement (Brown and Wilson 1956; Blair 1974; Butlin and Ritchie 1994; Servedio and Noor 2003; Pfennig and Pfennig 2009) and reinforcement of reproductive isolation. An interesting question that arises then is whether differences in reproductive traits correlate in recently diverged species, for instance, do differences in reproductive morphology correlate with differences in reproductive behaviour? And are these differences sufficiently large to function as prezygotic reproductive barriers, leading to reproductive isolation? Under a scenario of reinforcement in sympatry, we might expect that divergent reproductive traits will serve as fairly effective reproductive barriers (though not all sympatric species will necessarily be completely reproductively isolated). In contrast, species that have speciated in allopatry may lack (complete) reproductive isolation due to incomplete pre- or postzygotic barriers, despite having diverged in their reproductive traits. Secondary contact between such species may then result in the production of viable and potentially even fertile hybrid offspring.

Even in the absence of successful hybridization, both heterospecific mating attempts and actual heterospecific matings can result in wastage of energy, resources, time and/or gametes. This can lead to reproductive interference, which is defined as heterospecific reproductive activities that reduce the fitness of at least one of the species involved (Gröning and Hochkirch 2008; Kyogoku 2015; Grether et al. 2017; Shuker and Burdfield-Steel 2017). Interestingly, reproductive interference may be asymmetric, in that the fitness of one species is affected to a greater extent than that of the other (Gröning and Hochkirch 2008).

In our study, we investigated reproductive barriers and reproductive interference in two species of the free-living flatworm genus *Macrostomum*, namely *M. lignano*, an established model for studying sexual reproduction in hermaphrodites (Ladurner et al. 2005), and the recently described *M. janickei*, the currently most closely related congener known (Schärer et al. n.d.). Specifically, we examined if differences in the stylet morphology between these species correlated with differences in their mating behaviour and if they had similar fecundity. Furthermore, we investigated the potential for hybridization between the two species, and tested whether the resulting hybrids were fertile. Next, using geometric morphometrics we compared the stylet morphology of the parental species and the hybrids. Finally, we performed a mate choice experiment to test if individuals preferentially mated with conspecifics over heterospecifics, since this form of assortative mating could serve as a premating barrier between these two closely related species in a putative zone of sympatry.

## Materials and Methods

### Study organisms

*Macrostomum lignano* Ladurner, Schärer, Salvenmoser and Rieger 2005 and *M. janickei* Schärer in press are free-living flatworm species (Macrostomorpha, Platyhelminthes) found in the upper intertidal meiofauna of the Mediterranean Sea (Schärer et al. n.d.; Ladurner et al. 2005; Zadesenets et al. 2016, 2017). Despite being very closely related sister species (Schärer et al. n.d.), the morphology of their stylet is clearly distinct (see Figure 4 and results). *M. lignano* has a stylet that is “slightly curved, its distal opening [having a] slight asymmetric thickening” (Ladurner et al. 2005), while *M. janickei* has a more complex stylet that is a “long and a gradually narrowing funnel that includes first a slight turn (of ∼40°) and then a sharp turn (of >90°) towards the distal end […], giving the stylet tip a hook-like appearance.” (Schärer et al. n.d.).

Previous studies have shown that *M. lignano* is an outcrossing, reciprocally copulating species with frequent mating (on average about 6 copulations per hour, Schärer et al. 2004). Specifically, reciprocal copulation consists of both partners mating in the male and female role simultaneously, with reciprocal insertion of the stylet into the female antrum (the sperm-receiving organ) of the partner, and transfer of ejaculate consisting of both sperm and seminal fluids. Copulation is then often followed by a facultative postcopulatory suck behaviour (Schärer et al. 2004, 2011; Vizoso et al. 2010), during which the worm bends onto itself and places its pharynx over its own female genital opening, while appearing to suck. This behaviour is thought to represent a female resistance trait that has evolved due to sexual conflict over the fate of received ejaculate. Specifically, it is likely aimed at removing ejaculate components from the antrum, and sperm is often seen sticking out of the antrum after a suck (Marie-Orleach et al., 2013; Schärer et al., 2011; Schärer et al., 2004; Vizoso et al., 2010).

The individuals of *M. lignano* used in this experiment were either from the outbred LS1 culture (Marie-Orleach et al. 2013) or from the transgenic outbred BAS1 culture, which was created by backcrossing the GFP-expressing inbred HUB1 line (Janicke et al. 2013; Marie-Orleach et al. 2014) onto the LS1 culture (Marie-Orleach et al. 2016), subsequently cleaned from a karyotype polymorphism that segregates in HUB1 (Zadesenets et al. 2016, 2017), and finally bred to be homozygous GFP-positive (Vellnow et al. 2018). The *M. janickei* worms used were from a culture that was established using individuals collected from Palavas-les-Flots, near Montpellier, France (Schärer et al. n.d.; Zadesenets et al. 2016, 2017). Both species are kept in mass cultures in the laboratory at 20 °C in glass Petri dishes containing either f/2 medium (Andersen et al. 2007) or 32‰ artificial sea water (ASW) and fed with the diatom *Nitzschia curvilineata*.

### Experimental design

#### Experiment 1: Reproductive behaviour and hybridization

On day 1, for each species, we distributed 240 adult worms over 4 petri dishes with algae and ASW (using the transgenic BAS1 culture for *M. lignano*). On day 4, we removed the adults, such that the eggs were laid over a 3-day period, and the age of the resulting hatchlings did not differ by more than 3 days. On day 9 (i.e. well before the worms reach sexual maturity), we isolated these hatchlings in 24-well tissue culture plates (TPP, Switzerland) in 1 ml of ASW with *ad libitum* algae. Starting on day 34 and spread over 3 subsequent days, we then examined the mating behaviour by pairing these previously isolated and by then adult worms (as judged by their visible testes and ovaries) in one of three pairing types, namely *M. lignano* pairs (*M. lignano* × *M. lignano*, n = 57), *M. janickei* pairs (*M. janickei* × *M. janickei*, n = 57), or heterospecific pairs (*M. lignano* × *M. janickei*, n = 57).

Each observation chamber (Schärer et al. 2004) was assembled by placing 9 mating pairs (3 pairs of each pairing type) in drops of 3 μl of ASW each between two siliconized microscope slides separated by 257 μm, for a total of 19 observation chambers (i.e. 7, 4, and 8 chambers on the three subsequent days, respectively). The observation chambers were filmed under transmitted light for 2h at 1 frame s^−1^ with digital video cameras (DFK 41AF02 or DFK 31BF03, The Imaging Source) in QuickTime format using BTV Pro 6.0b7 (http://www.bensoftware.com/), and the resulting movies were scored manually frame-by-frame using QuickTime player. We used two different movie setups for filming the mating and they differed slightly in the cameras and light sources used.

After the two-hour mating period, we isolated both individuals of the heterospecific pairs, and one randomly chosen individual each of the *M. lignano* and *M. janickei* pairs, respectively, in 24-well plates and subsequently transferred them weekly to new plates. To obtain an estimate of the (female) fecundity resulting from these pairings the offspring production of these maternal individuals was followed and counted for 14 days (since worms eventually run out of stored sperm, Janicke et al. 2011). For each heterospecific pair, the number of (hybrid F1) offspring produced was averaged over both maternal individuals. And by confirming that all maternal offspring of the GFP-negative *M. janickei* were GFP-positive, we could ascertain that the GFP-positive BAS1 *M. lignano* had indeed sired these F1 hybrids. Moreover, previous experiments had shown that neither species self-fertilizes over a comparable observation period (Schärer and Ladurner 2003; Singh et al. 2019), thus any offspring produced in the heterospecific pairs must have resulted from outcrossing with the partners.

For each mating pair, we scored the movie up to the fifth copulation and observed the following copulation traits: copulation latency (i.e. time to first copulation), copulation duration, copulation interval, time of suck (after copulation), suck duration, and the number of sucks, while being blind with respect to both the pairing type and the species identity of individuals in the heterospecific pairs (note that the GFP-status of a worm cannot be determined under normal transmitted light). The decision to observe the behaviour up to and including the fifth copulation was made *a priori* (see also Marie-Orleach et al. 2013), and was motivated by our desire to get accurate estimates for each behaviour, by averaging all traits (except copulation latency) over this period for each pair and to keep the total observation time manageable. The copulation behaviour was defined as in Schärer et al. (2004), and the copulatory duration was measured starting from the frame when the pair was first tightly interlinked (like two small interlocking G’s) with the tail plates in close ventral contact, to the frame where their tail plates were no longer attached to each other. We scored a behaviour as a copulation only if the pair was in this interlinked position for at least 5 seconds. The copulation interval was measured as the duration between the end of a copulation to the start of the next copulation. The time of suck was measured (for sucks that followed a copulation, observed up to the fifth copulation) as the time elapsed between the end of the copulation preceding the suck and the start of the suck in question. The suck duration was measured from the frame where the pharynx was placed on the female genital opening up to the frame where the pharynx disengaged. The number of sucks was measured as the number of sucks observed up to the fifth copulation. The copulation duration, copulation interval, time of suck, and suck duration was averaged over all occurrences in a replicate.

The final sample sizes varied for the different behavioural traits, depending on how many replicates exhibited the particular trait of interest. We, respectively, excluded 3, 7 and 2 replicates of the *M. lignano* pairs, heterospecific pairs and *M. janickei* pairs from all analyses, since these replicates showed no copulations. In addition, 3 replicates of *M. janickei* had only one copulation, so we could not calculate the copulation interval for these drops. Moreover, in some replicates there were no sucks, which reduced our sample size for the time of suck and suck duration. The suck is considered a postcopulatory behaviour, and we therefore might not expect an individual to exhibit the postcopulatory behaviour unless it copulates. Thus, to examine if the number of sucks differed between the pairing types, we considered only the subset of drops in which we observed at least five copulations. Additionally, for offspring number we lost 2 replicates each for the *M. lignano* and *M. janickei* pairs. The final sample sizes are given in the respective figures.

#### Experiment 2: Hybrid fertility

We assessed the fertility of the F1 hybrid offspring from experiment 1, by pairing for 7 days a subset of the virgin hybrids with, respectively, virgin adult *M. lignano* (n = 24) or virgin adult *M. janickei* (n = 24) partners (grown up under identical conditions as the parents, but using the wildtype LS1 culture for *M. lignano*) and then isolating both the hybrids and their partners for 14 days to determine offspring number. By confirming that at least some of the F2 offspring from the crosses between the GFP-heterozygote F1 hybrids and the GFP-negative parents were GFP-positive, we could ascertain that we were indeed seeing successful backcrosses. We did not statistically analyse if offspring number differed depending on which parental species the hybrid was backcrossed onto, as the hybrids used were not statistically independent (e.g. some of them were siblings). Thus, we only descriptively examined offspring number produced from the backcrossing.

#### Experiment 3: Hybrid and parental species stylet morphology

To investigate the stylet morphology of the F1 hybrids, we compared the stylets of isolated virgin hybrids (n = 29; measured before the backcrossing experiment), to those of isolated *M. lignano* (n=25, from Ramm et al. 2019) and *M. janickei* (n=18, from Singh et al. 2019), using a geometric morphometrics landmark-based method (Zelditch et al. 2004). Briefly, worms were relaxed using a solution of MgCl_2_ and ASW, and dorsoventrally squeezed between a glass slide and a haemocytometer cover glass using standardised spacers (40 µm). Stylet images were then obtained at 400x magnification (Figure 4a-c), with a DM 2500 microscope (Leica Microsystems, Heerbrugg, Switzerland) using a digital camera (DFK41BF02, The Imaging Source, Bremen, Germany) connected to a computer running BTV Pro 6.0b7 (Ben Software). For geometric morphometrics, we placed a total of 60 landmarks on each stylet, two fixed landmarks each on the tip and base of the stylet and 28 equally spaced sliding semi-landmarks each along the two curved sides of the stylet between the base and the tip (Figure 4d-f), using tpsDig 2.31 (F. James Rohlf, 2006, Department of Ecology and Evolution, SUNY, http://life.bio.sunysb.edu/morph/), while being blind to the identity of the individual. Note that this landmark placement differs somewhat from that used earlier in *M. lignano* (Janicke and Schärer 2009) on account of the different morphology of the *M. janickei* stylet. Specifically, landmarks should represent homologous points on a morphological structure, and we here defined only four fixed landmarks that could be recognised in the F1 hybrids and both parental species (compared to six in *M. lignano* earlier), while more sliding semi-landmarks were used here to approximate the considerably more complex shape of the *M. janickei* stylet (i.e. 56 semi-landmarks now vs. 18 in *M. lignano* earlier). We always placed landmarks 1-30 on the stylet side that was further from the seminal vesicle (the sperm storage organ located near the stylet), while landmarks 31-60 were placed on the stylet side that was closer to the seminal vesicle (see Figure 4d-f). Also, to ensure that the orientation of the seminal vesicle and stylet with respect to the viewer was similar across all images, we mirrored the images for some specimens. We used tpsRelw 1.70 (http://life.bio.sunysb.edu/morph/) to analyse the resulting landmark configurations and extract the centroid size (an estimate of the size of the landmark configuration that can serve as a measure of the stylet size) and the relative warp scores (which decompose the total shape variation into major axes of shape variation). Our analysis yielded 71 relative warp scores, of which the first three relative warp scores explained 88% of all variation in stylet shape. For our statistical analysis, we here only focus on the first relative warp score (RWS1), as it explained 64% of the shape variation and captured the most drastic change in the stylet shape, including the extent of the stylet tip curvature (Figure 4g-i).

#### Experiment 4: Mate preference experiment

We assessed the mate preferences of *M. lignano* (BAS1) and *M. janickei* by joining two individuals of each species in 3 μl drops of ASW (for a total of 4 individuals per drop). In each of the four drops per observation chamber, the individuals of either one or the other species were dyed in order to permit distinguishing the species visually in the movies (i.e. *M. lignano* or *M. janickei* were dyed in two drops each per mating chamber). We dyed the worms by exposing them to a solution of the food colour Patent Blue V (Werner Schweizer AG, Switzerland, at 0.25 mg/ml of 32‰ ASW) for 24h. Patent Blue V does not affect the mating rate of *M. lignano* (Marie-Orleach et al. 2013), or of *M. janickei*, as the mating rate of dyed and undyed worms was similar (see Supplementary Figure S1).

In total, we constructed 17 observation chambers and filmed them under transmitted light for 2h at 1 frame s^−1^ (as outlined above), and the resulting movies were scored manually frame-by-frame using QuickTime player, while being blind to which species was dyed. For each drop, we determined the copulation type of the first copulation, i.e. conspecific *M. lignano*, conspecific *M. janickei* or heterospecific (*M. lignano* × *M. janickei*), and we also estimated the copulation frequencies of the three copulation types over the entire 2h period.

Out of the total 68 filmed drops we had to exclude 9 drops, 5 of which had an injured worm and 4 of which (one entire observation chamber) had dim lighting that made it difficult to distinguish the dyed worms. Thus, our final sample size was 59 drops.

### Statistical Analyses

In experiment 1, we constructed one-way ANOVAs with the pairing type (*M. lignano* pairs, heterospecific pairs, and *M. janickei* pairs) as the independent fixed factor, and using copulation latency, average copulation duration, average copulation interval, average time of suck, and average suck duration as the dependent variables, followed by post-hoc comparisons between the pairing types using Tukey’s honest significant difference (HSD) tests. Note that all conclusions remained unchanged if the two movie setups were included as a factor (data not shown). Data was visually checked for normality and homoscedasticity and log-transformed for all the above variables. For average time of suck, however, we added 1 to each data point before log-transformation, to avoid infinite values, since some sucks began immediately after copulation, leading to zero values. For the number of sucks and the offspring number we used Kruskal-Wallis tests (since these data could not be appropriately transformed to fulfil the assumptions for parametric tests), followed by post-hoc tests using Mann–Whitney–Wilcoxon tests with Bonferroni correction. Moreover, for all behaviours we calculated the coefficient of variation (CV) to evaluate how stereotypic the behaviour is for each pairing type. For all behaviours (except for the number of sucks), we calculated the CV for log-transformed data using the formula 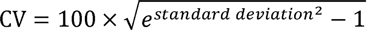 (Canchola 2017), while for number of sucks we calculated the CV for raw data using 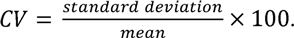

In experiment 3, we constructed one-way ANOVAs with the types of worm (*M. lignano*, *M. janickei*, or hybrid) as the independent fixed factor, and the centroid size and RWS1 as the dependent variables, followed by post-hoc comparisons using Tukey’s HSD. Note that these analyses need to be interpreted with some care, since the three groups we compared were not grown and imaged as part of the same experiment (though using the same methodology).

In experiment 4, three different copulation types could occur (i.e. *M. lignano* conspecific, heterospecific, and *M. janickei* conspecific), and to generate a null hypothesis of the expected proportions of each copulation type, we initially assumed random mating and hence no mating preference for either conspecific or heterospecific individuals in either species. Thus, under these assumptions the null hypothesis for the expected proportion of drops having these different copulation types as the first copulation was: *M. lignano* conspecific : heterospecific : *M. janickei* conspecific = 0.25 : 0.50 : 0.25. For each copulation type, we then determined the observed proportion of drops in which it was the first copulation, and examined if these proportions differed significantly from this null hypothesis, using a Chi-square goodness-of-fit test.

Next, we looked at the observed proportion of the three copulation types within each drop and across all drops, and as the null hypothesis we again used the same expected proportions as above. To test if the observed proportion of the three copulation types differed from this null hypothesis, we used repeated G-tests of goodness-of-fit (McDonald 2014), an approach that involves sequential tests of up to four different hypotheses, which, depending on the obtained results, will not all necessarily be carried out. The first hypothesis tests if the observed proportions within each drop fit the expectations. The second hypothesis examines if the relative observed proportions are the same across all drops by calculating a heterogeneity value. The third hypothesis examines if the observed proportion matches the expectation when the data is pooled across all drops. And finally, the fourth hypothesis examines if overall, the data from the individual drops fit our expectations using the sum of individual G-values for each replicate (obtained from testing the first hypothesis). Following this approach, we first calculated a G-test goodness-of-fit (with Bonferroni correction) for each drop. Second, this was followed by a G-test of independence on the data in order to obtain a ‘heterogeneity G-value’, which permits to evaluate if the drops differ significantly from each other. Since, this test revealed significant heterogeneity between the drops (see results), we did not pool the data or proceed with the remaining two tests, but instead drew our conclusion from the above G-tests of goodness-of-fit (corrected for multiple testing).

As we show in the results, in most drops, the majority of copulations were of the *M*. *lignano* conspecific type, followed by the heterospecific type (Figure 6a). To check whether this could be due to an intrinsically higher mating rate of *M. lignano* (see results), we generated a new null hypothesis that takes the observed mating rates of both *M. lignano* and *M. janickei* into account. For each drop, we therefore first calculated the mating rate of *M. lignano* as

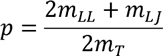

and similarly, the mating rate of *M. janickei* as

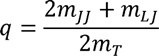

Where, *m_LL_*, *m_LJ_*, and *m_JJ_*, represent the observed numbers of *M. lignano* conspecific, heterospecific, and *M. janickei* conspecific copulations, and *m_T_* represents the total number of copulations (i.e. summed across all copulation types). Thus, we obtained a *p* and *q* value for each drop and if both species had the same mating rate, then we would expect *p* = *q* = 0.5. However, the results of the above analysis showed that *M. lignano* and *M. janickei* differed greatly in their mating rates (Figure 6b).

We, for each drop, therefore calculated the expected numbers of the different copulation types, given the observed mating rates *p* and *q* as

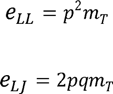

and

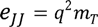

respectively, where *e_LL_*, *e_LJ_*, and *e_JJ_*, represent the expected numbers of *M. lignano* conspecific, heterospecific, and *M. janickei* conspecific copulations. Using these we then tested whether the resulting expected proportions were significantly different from the observed proportions for each drop, using a Chi-square goodness-of-fit test with Bonferroni correction for multiple testing. This allowed us to examine if the apparent preference of *M. lignano* for mating with conspecifics (i.e. the observed assortative mating) simply stemmed from the mating rate differences between the species, as opposed to a more explicit preference for conspecific partners.

All statistical analyses were carried out in R, version 3.1.1 (R Development Core Team, 2016).

### Ethical note

All animal experimentation was carried out in accordance to Swiss legal and ethical standards.

## Results

### Experiment 1: Reproductive behaviour and hybridization

The three pairing types differed in their mating behaviour, though to varying degrees for the different copulation traits. Pairing type had a significant effect on copulation latency (F_2,156_ = 4.688, P = 0.01; Figure 1a), with *M. lignano* pairs starting to copulate earlier than heterospecific pairs, while the *M. janickei* pairs had an intermediate copulation latency. The pairing type also had a significant effect on the copulation duration (F_2,156_ = 370.6, P < 0.001; Figure 1b), with *M. janickei* pairs having a nearly five-fold higher copulation duration than *M. lignano* pairs and heterospecific pairs, which did not significantly differ amongst themselves. Moreover, the copulation interval was affected by the pairing type (F_2,153_ = 8.124, P < 0.001; Figure 1c). *M. janickei* pairs had a significantly longer interval between copulations than *M. lignano* pairs, while the heterospecific pairs had intermediate copulation interval.

**Figure 1.**
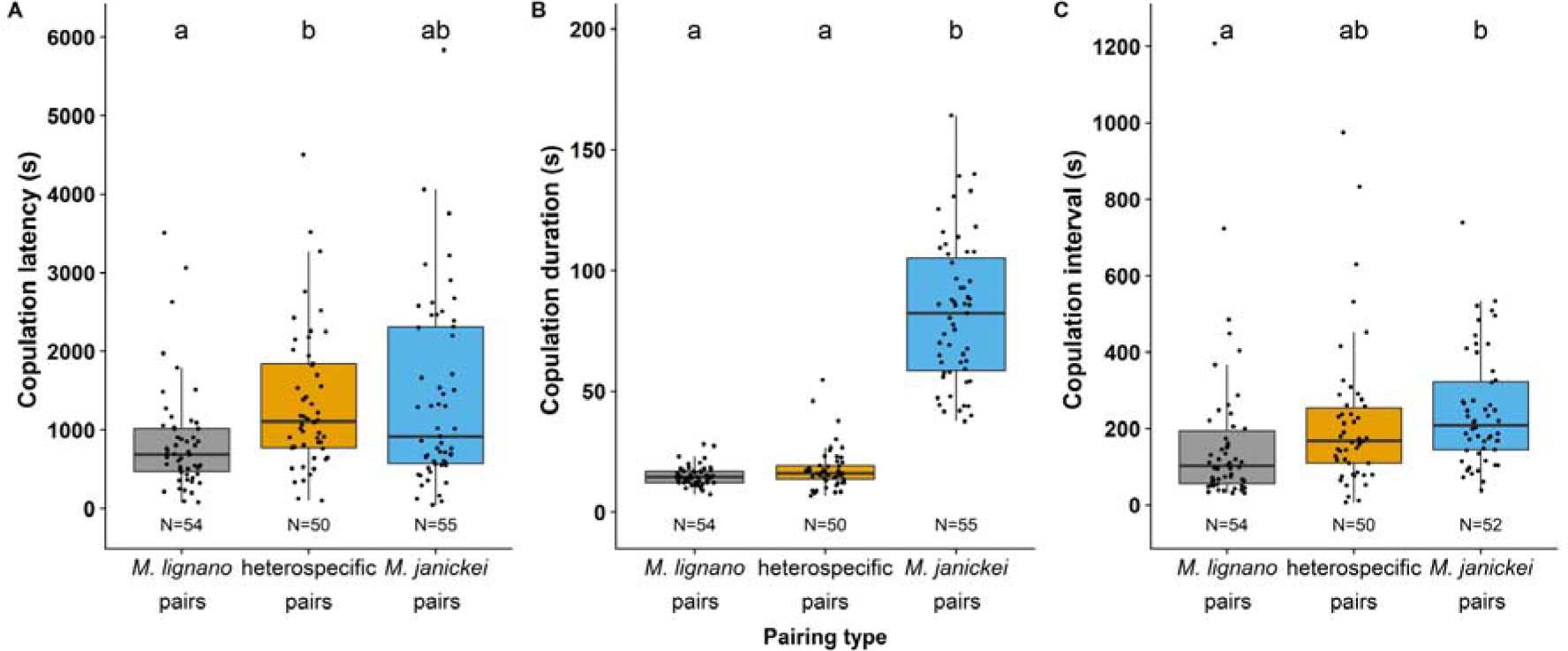
Boxplots of the a) copulation latency, b) (average) copulation duration, and c) (average) copulation interval of the three pairing types. Different letters denote significantly different effects inferred from Tukey HSD post-hoc tests. The boxplots display the 25th percentile, median, and 75th percentile and the whiskers represent the 5th and the 95th percentiles of the raw data, but note that log-transformed data was used for statistical analysis of all variables shown here. Sample sizes are given at the bottom of the plots.

For the suck behaviour, very few heterospecific replicates exhibited the behaviour, leading to a reduction in our sample size for the time of suck and suck duration (Figure 2). The time of suck (after copulation) differed between the pairing types (F_2,92_ = 48.15, P < 0.001; Figure 2a), with *M. lignano* pairs usually sucking almost immediately after copulation, while the *M. janickei* pairs and heterospecific pairs took a longer time to start sucking. The suck duration was also significantly affected by the pairing type (F_2,92_ =7.80, P < 0.001; Figure 2b), with *M. janickei* pairs having a longer suck duration than *M. lignano* pairs, while the heterospecific pairs did not significantly differ from the other two pairing types. Interestingly, the number of sucks was significantly affected by the pairing type (Kruskal–Wallis test: χ^2^ = 41.16, df = 2, P < 0.001; Figure 2c), with *M. lignano* pairs sucking most frequently, followed by the *M. janickei* pairs. The heterospecific pairs sucked least frequently.

**Figure 2.**
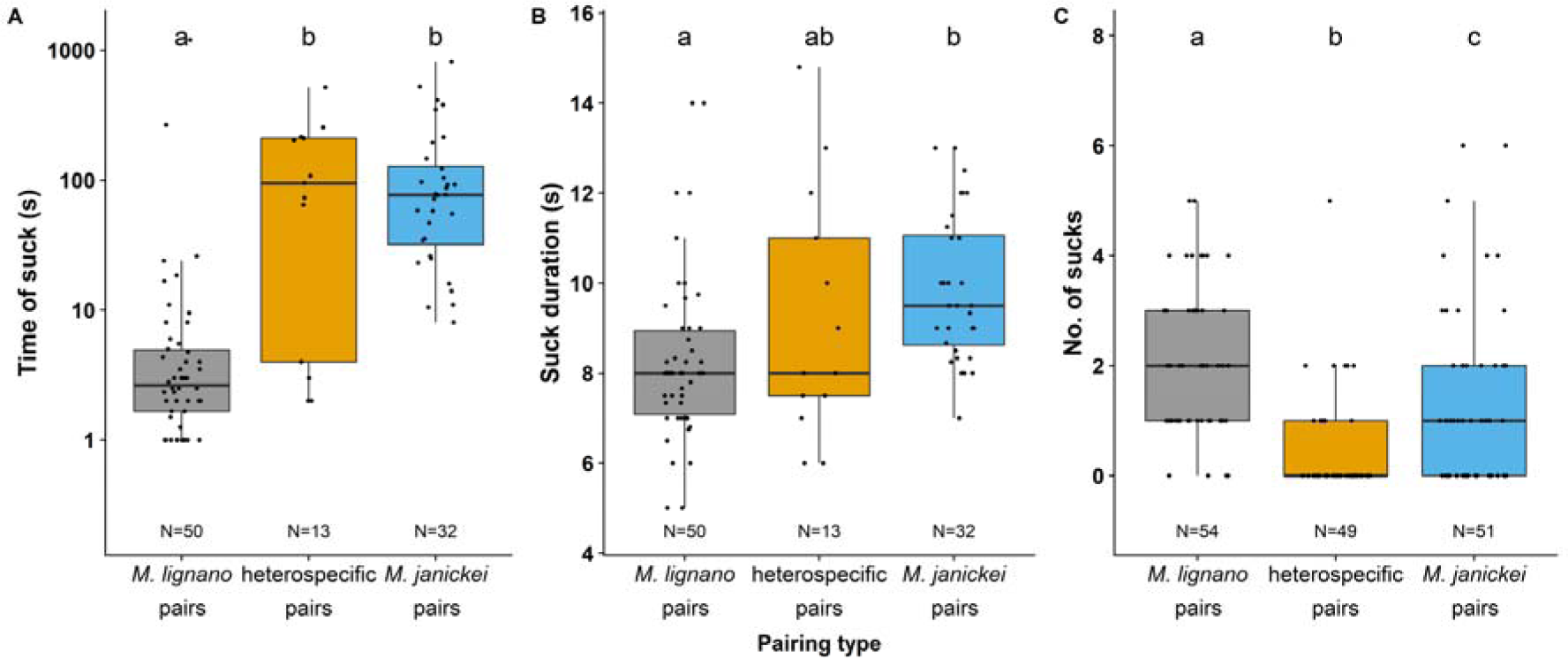
Boxplots of the a) (average) time of suck (after copulation), b) (average) suck duration, and c) number of sucks of the three pairing types (recall that we here only consider pairs that copulated at least 5 times). Different letters denote significantly different effects inferred from Tukey HSD post-hoc tests (for a and b) or Mann–Whitney–Wilcoxon tests with Bonferroni correction (for c). The boxplots display the 25th percentile, median, and 75th percentile and the whiskers represent the 5th and the 95th percentiles of the log-transformed data (for a) and the raw data for (b and c), but note that log-transformed data was used for statistical analysis (for a and b). We added 1 to each data point for time of suck before log-transforming to avoid infinite values (see text for details). Sample sizes are given at the bottom of the plots.

Remarkably, for most behaviours the heterospecific pairs had the highest CV, suggesting that heterospecific behaviour was relatively variable and less stereotypic than conspecific behaviour (Table 1).

**Table 1.** The coefficient of variation (CV) of each pairing type for all behaviours. For most behaviours the heterospecific pairs had the highest CV.

| behaviour | *M. lignano* pairs | heterospecific pairs | *M. janickei* pairs |
| --- | --- | --- | --- |
| copulation latency | 86 | 88 | 127 |
| copulation duration | 27 | 44 | 39 |
| copulation interval | 100 | 116 | 69 |
| time of suck | 234 | 810 | 175 |
| suck duration | 21 | 29 | 16 |
| No. of sucks | 66 | 209 | 120 |

In addition, while heterospecific pairs were capable of producing hybrid offspring—a new finding for this genus—they produced significantly fewer offspring than conspecific pairs (Kruskal-Wallis test: χ^2^ = 48.04, df = 2, P < 0.001; Figure 3a), which had a comparable fecundity. Out of the 10 heterospecific replicates that produced hybrids, in 6 replicates only the *M. lignano* parent produced hybrids while in the other 4 replicates only the *M. janickei* parent produced offspring. Thus, hybridization was symmetrical, with each species being capable of inseminating and fertilizing the other.

**Figure 3.**
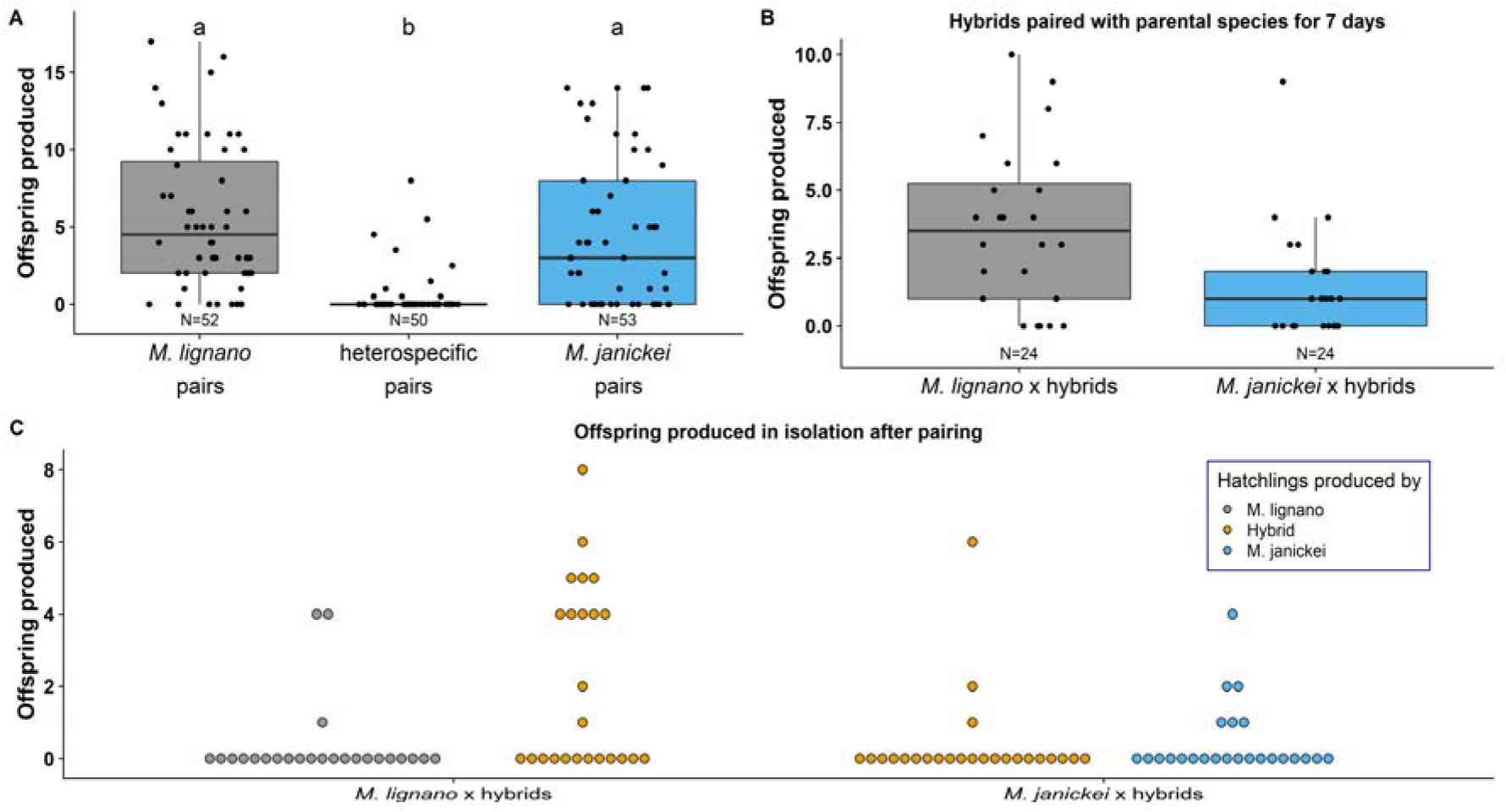
Plot of F2 hybrid offspring produced (female fecundity) a) by the three pairing types in Experiment 1; b) in the wells where the F1 hybrids were paired with an individual of one of their parental species for 7 days, and c) post-pairing isolated hybrid and parental individuals in Experiment 2. The boxplots in a) and b) display the 25th percentile, median, and 75th percentile and the whiskers represent the 5th and the 95th percentiles of the raw data, while c) is a dotplot. Note that in c) each backcrossed pair is represented twice as each pair comprises a hybrid and a parental species individual, so the replicates are not independent and the figure is only for visualisation. Sample sizes are given at the bottom of the plots in a) and b).

**Figure 4.**
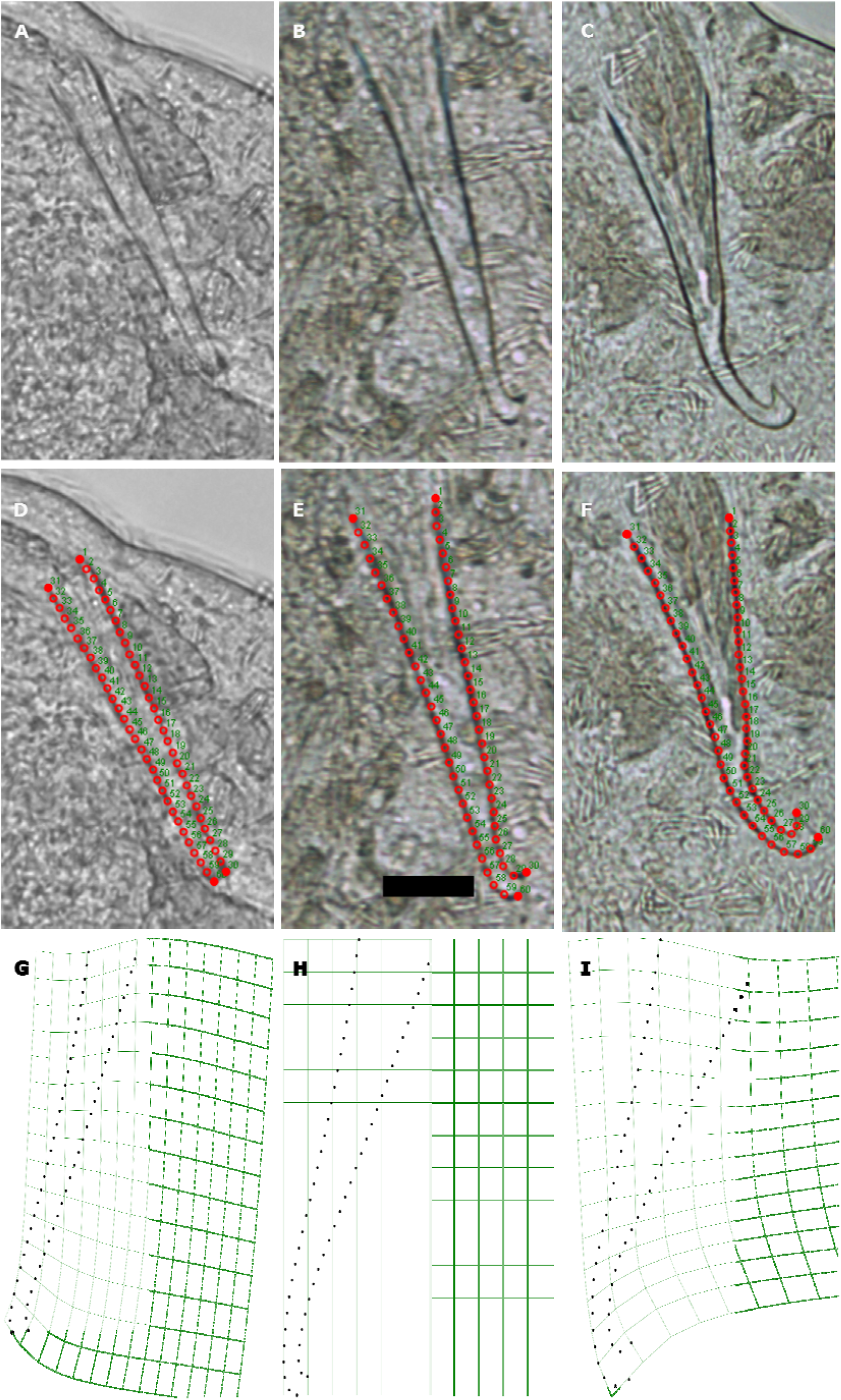
Morphology and geometric morphometrics of the stylet. Micrographs of the stylet of an individual a) *M. lignano*, b) F1 hybrid, and c) *M. janickei*. The placement of 60 landmarks along the stylet of an individual d) *M. lignano*, e) F1 hybrid and f) *M. janickei*. Note that we placed four fixed landmarks (filled red circles), two on the stylet base and two on the stylet tip, and 28 equally spaced sliding semi-landmarks (empty red circles) along each curved side of the stylet. The numbers indicate the order in which the landmarks were placed (note that the seminal vesicles always are to the left of the stylet). Visualization of thin-plate splines of the stylet derived from relative warp score analysis. Each panel shows the visualization for the mean relative warp score 1 (RWS1) of individuals of g) *M. lignano*, h) the F1 hybrids and i) *M. janickei*. Thus, in general *M. lignano* has a relatively straight stylet tip and *M. janickei* has a stylet tip that curves drastically, while the hybrids have intermediate curvature. The scale bar in e) represents 20 µm, and is applicable to all photomicrographic images.

### Experiment 2: Hybrid fertility

Most of the F1 hybrids were fertile and produced offspring in the wells while paired with worms from the parental species. Specifically, we found that 19/24 and 14/24 pairs of *M. lignano* × hybrid and *M. janickei* × hybrid produced hybrid F2 offspring, respectively, while they were paired with an individual of one of their parental species for 7 days (Figure 3b), while post-pairing, relatively few individuals of either hybrids or parentals produced offspring in isolation (Figure 3c).

### Experiment 3: Hybrid and parental species stylet morphology

The stylet morphology was significantly different between *M. lignano*, *M. janickei* and the F1 hybrids (Figure 4). The centroid size, an estimate of stylet size, was different between the groups (F_2,69_ = 33.26, P < 0.001; Figure 5a), with the F1 hybrids having a larger centroid size than *M. lignano* and *M. janickei*, which did not differ amongst themselves. The RWS1 of the stylets, which primarily seemed to capture variation in the curvature of the stylet tip and the width of the stylet base (Figure 4g-i), was significantly different between all groups (F_2,69_ = 238, P < 0.001; Figure 5b), with the RWS1 of the hybrids being intermediate between that of *M. lignano* and *M. janickei*, indicating that the shape of hybrid stylet was morphologically intermediate between the parental species.

**Figure 5.**
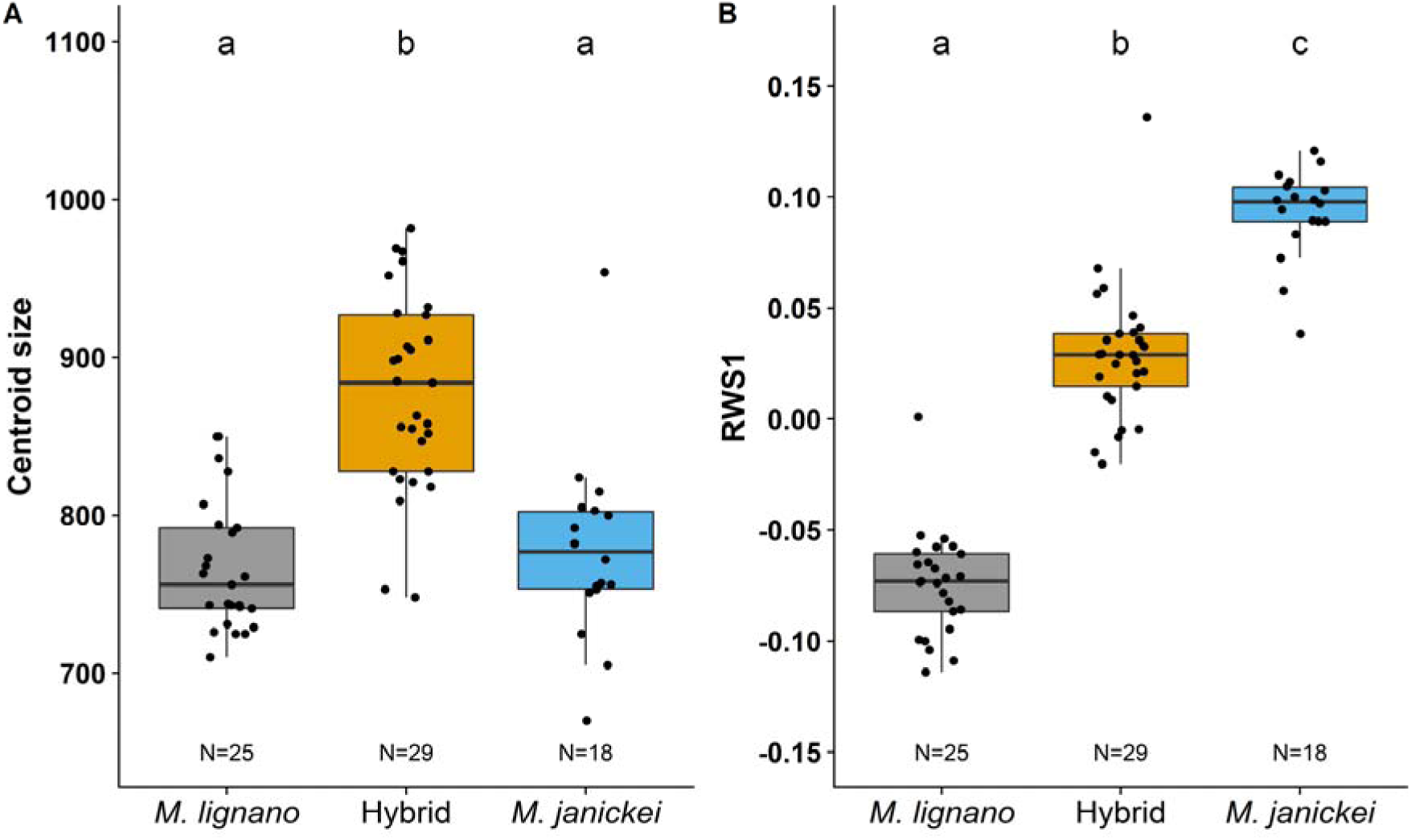
Boxplot for a) centroid size and b) relative warp score 1 (RWS1) of the stylets of *M. lignano*, F1 hybrid and *M. janickei* worms. Different letters denote significantly different effects inferred from Tukey HSD post-hoc tests. The boxplots display the 25th percentile, median, and 75th percentile and the whiskers represent the 5th and the 95th percentiles of the raw data. Sample sizes are given at the bottom of the plots.

**Figure 6.**
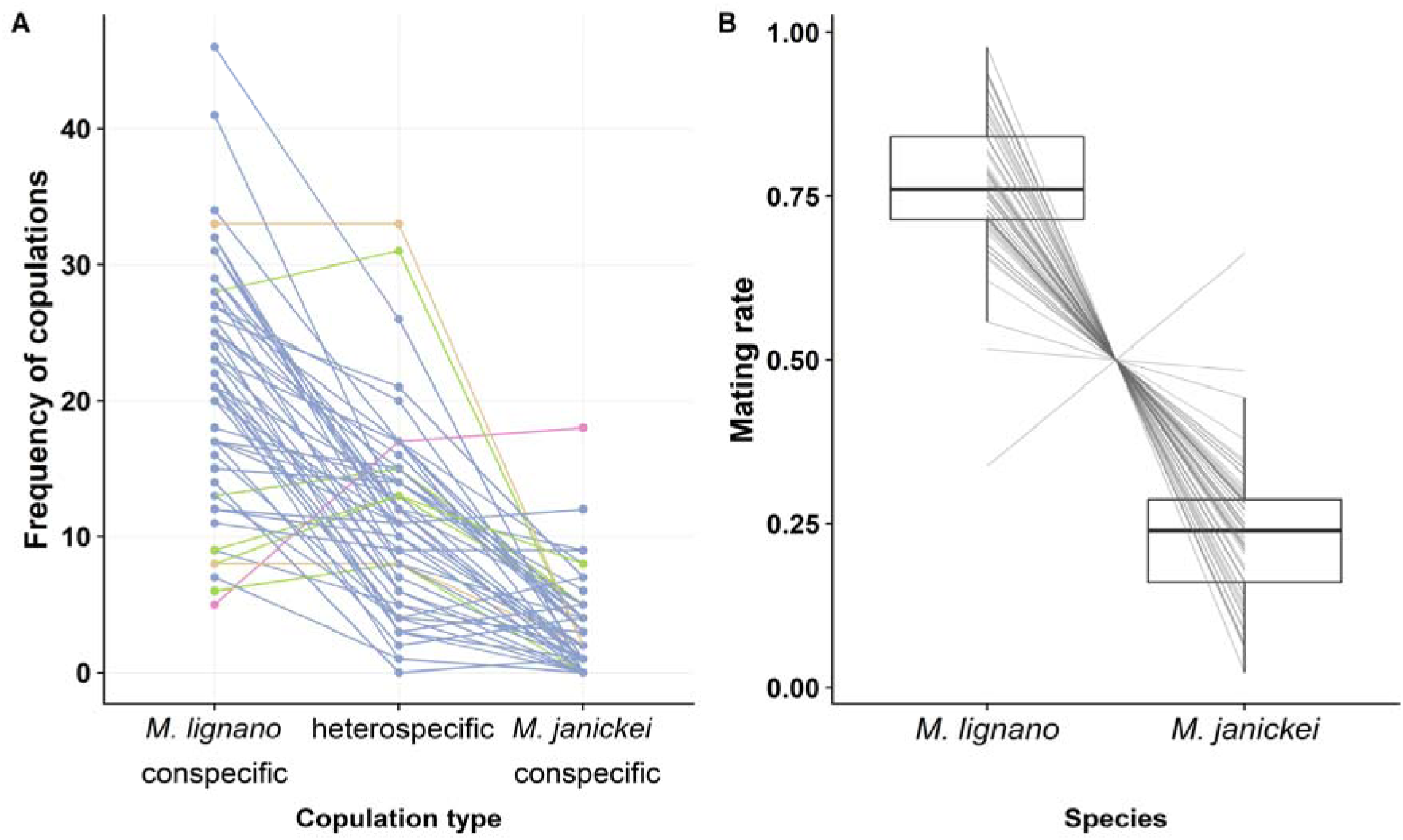
a) Frequency of *M. lignano* conspecific, heterospecific, and *M. janickei* conspecific copulations. Each line connects values obtained from the same drop. The different colours help to visualise which copulation type had the highest frequency in a drop (blue, *M. lignano* conspecific; green, heterospecific; pink, *M. janickei* conspecific; orange, same in *M. lignano* conspecific and heterospecific), b) Boxplot of mating rate of *M. lignano* and *M. janickei*. The boxplots display the 25th percentile, median, and 75th percentile and the whiskers represent the 5th and the 95th percentiles of the raw data. Each line connects values obtained from the same drop.

### Experiment 4: Mate preference experiment

Out of the 59 analysed drops, we found that 34 (57.6%) drops had a *M. lignano* conspecific copulation as the first copulation, while that was true for only 18 (30.5%) and 7 (11.9%) drops for heterospecifics and *M. janickei* conspecifics, respectively. These proportions differed significantly from our null hypothesis under random mating (Chi-square goodness-of-fit test: χ^2^ = 33.68, df = 2, P < 0.001).

With respect to the observed proportion of the different copulation types within drops, the data from 55 of the 59 drops (without Bonferroni-correction P < 0.05, Supplementary Table S2) differed significantly from the null hypothesis, though after Bonferroni correction that number dropped to just 46 drops (Bonferroni-corrected P < 0.05, Supplementary Table S2). Interestingly, we found significant variation in the observed proportion between the drops (‘heterogeneity G-value’ = 358.55, df = 116, P < 0.001), as is also evident from Figure 6a. The general trend was that *M. lignano* conspecific copulations were the most frequent, followed by heterospecific copulations, while we observed relatively few *M. janickei* conspecific copulations in most of the drops. In 51 drops, the *M. lignano* conspecific copulations were the most frequent, while in only one drop was the proportion of *M. janickei* conspecific copulations the highest (see colours in Figure 6a). Moreover, in five drops, the highest proportion of copulations was of the heterospecific type, while in two drops, *M. lignano* conspecific and heterospecific copulations jointly had the highest proportion. Surprisingly, we found that in 52 drops there was a higher proportion of heterospecific copulations than of *M. janickei* conspecific copulations (with zero *M. janickei* conspecific copulations in 13 drops), indicating that under these conditions, the *M. janickei* worms mated more often with a *M. lignano* heterospecific than with a *M. janickei* conspecific individual. This could either represent a preference in *M. janickei* for mating with *M. lignano*, or it could potentially also result from *M. lignano* having an intrinsically higher mating rate, which we explore next.

In our mate preference assays, the mating rate of *M. lignano* and *M. janickei* was indeed different, with *M. lignano* having a much higher mating rate than *M. janickei* (Figure 6b). When we took the mating rate differences between the two species into account, the Chi-Square goodness-of-fit test showed that in 55 out of 59 drops the observed and expected copulation frequencies were not significantly different (Bonferroni-corrected P > 0.05, Supplementary Table S3). This suggests that the difference in the copulation frequencies of the different copulation types, including the high frequency of heterospecific copulations in *M. janickei*, is largely explained by the intrinsic differences in mating rate of the two species, rather than stemming from an explicit preference for heterospecific partners.

## Discussion

Our study shows that the closely related species *M. lignano* and *M. janickei* differ significantly, not only in their stylet morphology, but also in several aspects of their mating behaviour. These considerable morphological and behavioural differences do not, however, appear to represent strong premating barriers, since the worms were readily able to engage in heterospecific matings. In contrast, there seem to be significant postmating barriers between these two species, as only few hybrid offspring were produced from these heterospecific matings. Moreover, the resulting hybrids were fertile, showing a stylet morphology that was intermediate between the parental species, and capable of backcrossing to both parental species. Interestingly, the data from our mate preference assay revealed distinct asymmetries in the mating patterns between the two species. While *M. lignano* clearly engaged predominantly in conspecific matings, thereby exhibiting assortative mating, *M. janickei* ended up mating more often with heterospecific individuals, and we suggest that both likely occurred as a result of the higher mating rate of *M. lignano* compared to *M. janickei*. In the following, we discuss these results in some more detail.

### Experiment 1: Reproductive behaviour and hybridization

A potential factor that could lead to the observed differences in behavioural traits between the two species is genital morphology. For example, a positive correlation between copulation duration and structural complexity of the intromittent organ has been reported in New World natricine snakes (King et al. 2009), wherein the authors hypothesized that the evolution of elaborate copulatory organ morphology is driven by sexual conflict over the duration of copulation. Similar to the findings of that study, the nearly five-fold longer copulation duration of *M. janickei* pairs compared to *M. lignano* pairs could in part be dictated by its considerably more complex stylet. Moreover, similar to the male genitalia, the female genitalia are also more complex in *M. janickei* than *M. lignano* (Schärer et al. n.d.). And in addition to copulation duration, the longer suck duration of *M. janickei* could also be correlated with the genital complexity, since removal of ejaculate from the more complex female genitalia might be more difficult and take more time.

In addition to genital morphology, both copulatory and post-copulatory behaviour might also be influenced by the quantity and composition of the ejaculate transferred during copulation. For example, a larger quantity of ejaculate might be accompanied by a longer copulation duration, and possibly also a longer suck duration, since the hypothesised function of the suck behaviour is to remove ejaculate components (Schärer et al. 2004; Vizoso et al. 2010). Moreover, a longer copulation duration might require longer phases of recovery during which spent ejaculate is replenished, leading to lower copulation frequency and a longer copulation interval. A previous study in *M. lignano* showed that pairs formed from virgin worms copulated approximately 1.6x longer than pairs formed from sexually-experienced worms, and also that individuals that had copulated with virgin partners had a lower suck frequency compared to individuals that had copulated with sexually-experienced partners (Marie-Orleach et al. 2013). This led the authors to hypothesize that virgin partners have more own sperm and seminal fluid available (which both were confirmed), and may thus transfer more ejaculate than sexually-experienced partners, and that some components of the ejaculate are aimed at manipulating the partner and preventing it from sucking (Marie-Orleach et al. 2013). Indeed, studies in *Drosophila* have shown the presence of non-sperm components in the ejaculate, which can alter the physiology, immunity, life history, and behaviour of the recipient, causing strong effects on the fitness of both the donor and the recipient (Chapman 2001; Perry et al. 2013; Schwenke et al. 2016; Billeter and Wolfner 2018). Efforts to elucidate the function of ejaculate components (like seminal-fluid proteins) in *M. lignano* have recently made considerable progress (Weber et al. 2018; Patlar et al. 2019; Ramm et al. 2019), and it will be interesting to see if these have similar functions.

Longer copulation intervals or temporal aspects of sucking (i.e. onset of sucking) could potentially also result from the action of some transferred ejaculate components that acts as a relaxant, leading to inactivity and delayed re-mating or delayed sucking. Interestingly, we noticed that very few individuals in the heterospecific pairs exhibited the suck behaviour, which could simply result from low or absent ejaculate transfer. It is also conceivable that sucking is triggered by species-specific ejaculate components and their interaction with the female reproductive organ, and hence the absence or low amounts of such components could result in fewer sucks. Alternatively, individuals of one species might be more effective at preventing suck in heterospecific partners, as heterospecific partners may lack coevolved defences against such ejaculate substances. Similar to our observation, a cross-reactivity study in the land snail, *Cornu aspersum*, showed that its diverticulum (a part of the female reproductive system) only responded to the love-dart mucus of some, but not other, land snail species, pointing towards species-specific effects of accessory gland products (Lodi and Koene 2016).

Moreover, the different behavioural components might be correlated with each other. For example, there could be a trade-off between the suck duration and suck frequency for ejaculate removal, such that longer sucks or more frequent sucks serve the same purpose. Similarly, a longer copulation duration might be accompanied by a longer suck duration and copulation interval (as discussed above). In support of this, we did see that *M. lignano* pairs had both a short copulation and suck duration, but a high copulation and suck frequency, while the converse was true for *M. janickei* pairs. Thus, there can be correlations between different aspects of reproductive behaviour and morphology, and a large-scale comparative study of reproductive behaviour and morphology in *Macrostomum* species would help to improve our understanding of the complexity and evolution of reproductive traits.

Heterospecific pairs showed higher CVs compared to the other two pairing types for both copulation duration and copulation interval, potentially suggesting disagreements over the optimal copulation duration and copulation frequency in these pairs. In addition, heterospecific pairs exhibited higher CVs compared to conspecific pairs for all suck related behaviours. Note that in these movies we could not visually distinguish the two species in the heterospecific pairs, but it appears likely that the short and immediate sucks were performed by *M. lignano* individuals, while the longer and delayed sucks were performed by *M. janickei* individuals. Interestingly, the suck behaviour seems to be a highly stereotypical behaviour, with the CVs being lower for suck duration than for copulation duration for each of the mating pair types. This is similar to what was noted from earlier behaviour studies of *M. lignano* (Schärer et al. 2004).

Whereas conspecific pairs of both species produced similar offspring numbers, heterospecific pairs gave rise to offspring relatively rarely, despite most pairs having copulated successfully, presumably due to postmating-prezygotic or postzygotic reproductive barriers. In our study, hybridization was symmetrical, with both species being able to inseminate and fertilize the other species. Interestingly, in none of the heterospecific replicates did both partners produce offspring. While this could point towards unilateral transfer of sperm during copulation, we cannot ascertain if this only occurs in heterospecific pairs or if conspecific pairs also show a similar pattern, as we collected only one partner for each conspecific pair. To the best of our knowledge this is the first study to have documented hybridization between species of the genus *Macrostomum*, and there is also very sparse information only about hybridization in free-living flatworms in general (Pala et al. 1982; Bullini 1985), while there is some more information about parasitic flatworms (Taylor 1970; Thèron 1989; Detwiler and Criscione 2010; Itagaki et al. 2011; Henrich et al. 2013).

### Experiment 2: Hybrid fertility

While historically, hybrids have often been considered to be sterile and evolutionary dead-ends (see Mallet 2005), hybridization sometimes leads to viable and fertile offspring. In such cases, hybridization can serve as a mechanism for generating diversification, by creating adaptive variation and functional novelty in morphology and genotypes (Mallet 2005; Bonnet et al. 2017), a view that has been reinforced by the widespread presence of allopolyploidy among plants (Soltis and Soltis 1995; Soltis et al. 2015; Wendel et al. 2016). In our study, heterospecific matings between *M. lignano* and *M. janickei* resulted in the production of viable hybrids, which we could successfully backcross onto both parental species. Though our study demonstrates hybridisation between the two species, we currently have no evidence for these species occurring in sympatry. *M. lignano* has previously been collected from locations in Greece and Italy, while *M. janickei* has to date only been collected from France (Schärer et al. n.d.; Zadesenets et al. 2016, 2017). Assuming this geographic distribution indicates absence of sympatric zones, it would follow that the observed reproductive trait divergence might not have occurred as a result of reinforcement of reproductive isolation. Thus, the differences in reproductive characters will not necessarily serve as reproductive barriers, and this could potentially explain our observed results.

Remarkably, both of our study species exhibit an unusual karyotype organization (Zadesenets et al. 2016), involving hidden tetraploidy and hexaploidy in *M. lignano* and *M. janickei*, respectively (likely as a result of a whole genome duplication event). Moreover, both species show additional chromosome number variation in the form of aneuploidies of the largest chromosome, also leading to other ploidy levels (Zadesenets et al. 2017). Interestingly, individuals with unusual karyotypes do not show behavioural or morphological abnormalities and reproduce successfully, at least in *M. lignano* (Zadesenets et al. 2016). The fact that we can obtain viable hybrids between the two species calls for studies of the resulting karyotypes of these F1 hybrids and the F2 backcrosses.

### Experiment 3: Hybrid and parental species stylet morphology

The parental species differed significantly in the morphology of their stylet, though their overall stylet size was similar. In contrast, the hybrids possessed a stylet that had a morphology that was intermediate between that of the parental species, but was distinctly larger in size, for which we currently have no explanation (as already mentioned above, these results need to be interpreted with some care, since the data used in this comparison stemmed from three separate experiments). A study in closely related species of damselflies had also shown that, despite differences in genitalia morphology, the species had incomplete mechanical isolation and could hybridize (Barnard et al. 2017). An interesting follow-up to our study would be to use QTL mapping in order to identify which gene regions are involved in stylet formation and shape (Tanaka et al. 2015; Fujisawa et al. 2019; Hagen et al. 2019), which would help us understand genital evolution (Yassin 2016). This approach might, however, be rendered difficult due to the karyotype polymorphisms present in the two *Macrostomum* species.

### Experiment 4: Mate preference experiment

Our mate preference experiment showed that there is some degree of assortative mating between *M. lignano* individuals, which appears to mostly stem from the higher mating rate of *M. lignano*. This is in line with our results from Experiment 1, where *M. lignano* conspecific pairs had shorter copulation latencies, shorter copulation durations and shorter copulation intervals compared to *M. janickei* conspecific pairs (Figure 1). Thus, mate choice in these two species seems to be governed mainly by behavioural characteristics, such as mating rate, rather than an explicit preference for a conspecific or heterospecific partner. A potential factor affecting mating rate could be sexual selection, for instance, in polygamous mating systems, sexual selection can select for persistent mating efforts, particularly in males, which in turn can lead to reproductive interference between the species (Gröning and Hochkirch 2008; Burdfield-Steel and Shuker 2011; Kyogoku 2015). Interestingly, a similar phenomenon has been observed in experimentally evolved populations of *Drosophila pseudoobscura* that experienced different sexual selection intensity regimes of either monogamy or polyandry (Snook et al. 2005; Debelle et al. 2014). A mate choice experiment showed that males from polyandrous populations had a higher probability of mating than those from monogamous populations (Debelle et al. 2016), potentially due to having evolved under strong male-male competition and hence initiating courtship faster and more frequently than monogamous males (Crudgington et al. 2010). Similarly, an experimental evolution study on a seed beetle, *Callosobruchus chinensis*, also showed that beetles evolved under a polygamous regime caused stronger reproductive interference on a congener species (*C. maculatus*) than beetles evolved under a monogamous regime (Kyogoku and Sota 2017; Kyogoku et al. 2019). In addition to the above examples, multiple empirical studies have proposed a role of sexual selection in occurrence of reproductive interference between species (Kyogoku and Sota 2015; Yassin and David 2016).

In our experiment, the over-representation of heterospecific matings in *M. janickei* could lead to asymmetric reproductive interference between these species. Though we did not explicitly investigate how fecundity is affected, it seems likely that *M. janickei* would pay a higher fitness cost compared to *M. lignano* in such a context. Future studies should explicitly investigate if and how mating rate differences can affect the fecundity of the species and whether the cost is symmetric for both species, or if *M. janickei* suffers more due to a reduced conspecific mating rate. Moreover, as we outlined above, while our study raises the interesting possibility of hybridization occurring in zones of secondary contact between the two species, we are currently not aware of any overlapping ranges of the two species (but this may largely be due to the lack of sampling effort). Considering their heterospecific interactions though, it might be difficult for the species to co-exist, since *M. lignano* might be expected to displace *M. janickei* from any overlapping zones due to potential asymmetric reproductive interference. Alternatively, selection for reinforcement of reproductive isolation might occur, leading to character displacement of the species in sympatric zones, such that heterospecific interactions are reduced.

### Conclusions

Our study shows that reproductive traits can evolve rapidly, even between closely related species, though they do not necessarily pose a reproductive barrier to hybridization. An interesting question that arises then is whether mating behaviour and genital morphology co-evolve or whether they diversify independently. A phylogenetic comparative study that looks at the evolution of these reproductive traits in more species across the *Macrostomum* genus would help us answer these open questions. Moreover, using hybridization and techniques like QTL mapping, we could aim at understanding the genetic basis of rapidly evolving and diversifying reproductive traits like mating behaviour and genitalia, and in turn broaden our understanding of speciation in free-living flatworms, a highly species-rich group of simultaneous hermaphrodites.

## Acknowledgements

We would like to thank Tim Janicke and Georgina Rivera Ingraham for help with collecting specimens of the study species *M. janickei*, Steven A. Ramm for providing access to the images of the *M. lignano* stylets from their study, Jeremias Brand for help with imaging the hybrid morphology, Nikolas Vellnow and Axel Wiberg for helpful discussions, and Gudrun Viktorin, Lukas Zimmerman, Jürgen Hottinger, Urs Stiefel and Daniel Lüscher for technical support. This research was supported by grant 31003A-162543 of the Swiss National Science Foundation (SNSF) to LS.

